# Brain-derived synaptic vesicles have an intrinsic ability to sequester tubulin

**DOI:** 10.1101/2025.05.08.652815

**Authors:** Tiago Mimoso, Aleksandr Korobeinikov, Alexander Stein, Dragomir Milovanovic, Silvio O. Rizzoli, Sarah Köster, Sofiia Reshetniak

## Abstract

The presence and function of microtubules within the synaptic bouton has long been under investigation. In recent years evidence has accumulated that connects the synaptic vesicle cluster to the local dynamics of microtubule ends. Nonetheless, one question remains open, namely whether the vesicles influence the availability of tubulin within the synaptic compartment. An analysis of previously published live imaging experiments indicates that tubulin is strongly enriched in the synaptic vesicle cluster. To analyze the vesicle-tubulin interaction directly, we isolated vesicles from the mouse brain and imaged them together with fluorescent tubulin *in vitro*. We found that soluble tubulin is collected by synaptic vesicles in physiological buffers, resulting in the formation of tubulin-rich regions (TRRs) on the respective vesicle clusters. We conclude that the synaptic vesicle cluster is indeed able to recruit soluble tubulin.

## Introduction

Synaptic transmission relies on the fusion of neurotransmitter-loaded synaptic vesicles (SVs) with the plasma membrane of the synaptic bouton, to deliver their contents into the synaptic cleft, and to thus activate postsynaptic receptors. The presence of SVs in the synaptic bouton is ensured by cytoskeleton-dependent transport of SV precursors, which are continually brought from the cell body, and are assembled into vesicles in the synapse (Chenouard *et al*., 2020; Petzoldt, 2023). Their interaction with microtubules and actin cytoskeleton continues throughout their lifetimes, as vesicles are continually transported between synapses (Darcy *et al*., 2006; Staras *et al*., 2010; Gramlich & Klyachko, 2017; Parkes *et al*., 2023) or even between synapses and the cell body (Wong *et al*., 2012). Recent works demonstrate that while the majority of SV employ actin cytoskeleton, a population of mature SVs rely on microtubules for their transport (Parkes *et al*., 2023), proposedly as they are targeted for trafficking back to the soma for degradation (Ivanova & Cousin, 2022).

While the transport-related interaction of SVs and microtubules is therefore well-established, with the presence of microtubules heavily influencing either the delivery of SV precursors to synapses or SV transport to the cell body, other effects of microtubules on SV dynamics are less clear. A detailed analysis of neurotransmission at a large central synapse, the calyx of Held, led to the observation that microtubules enter deep into the synapse, where they colocalize with a large number of resident (not transported) SVs (Babu *et al*., 2020), which fits well with many previous analyses of synapses by electron microscopy (e.g. (Gray *et al*., 1982; Gray & Katz, 1997). Moreover, microtubule depolymerization slowed down the recovery of the synapses from short-term depression (Babu *et al*., 2020), suggesting that such microtubules exhibit a functional interaction with SV clusters.

In principle, functional interactions between microtubules and SVs could take many other additional forms. The protein tau, well known from the context of diseases termed tauopathies (including Alzheimer’s Disease), is mainly considered to be a stabilizer of microtubules (Barbier *et al*., 2019). Tau is recruited to synapses, at least in pathological conditions, by binding to the SV membrane proteins synaptogyrin-3 (McInnes *et al*., 2018), where it seems to alter actin and microtubule assembly, by causing a local *de novo* polymerization, with deleterious effects on synaptic function (Zhou *et al*., 2017; Hori *et al*., 2022). Notably, the effects of tau mutations on synaptic transmission differ between animal models (Taipala *et al*., 2022), and tubulin binding adds another complicating factor to be considered in this area.

Other SV proteins may interact with tubulin, including highly abundant molecules as the calcium sensor synaptotagmin I, whose calcium-binding domains bind directly to the C-terminal region of beta-tubulin, and can cause tubulin polymerization *in vitro* (Honda *et al*., 2002). Two soluble synaptic proteins, which associate closely to SVs, synapsin I and alpha-synuclein, also bind microtubules, with the first being able to bring microtubules together, in bundles (Baines & Bennett, 1986), while the latter appears to increase the growth rate of microtubules, as well as the frequency of rapid shrinkage period (also termed microtubule catastrophe events; (Cartelli *et al*., 2016)).

Overall, these observations suggest that SVs may interact with tubulin, causing the retention of soluble tubulin within the presynaptic space, and thereby modulating the local microtubule dynamics. This is further suggested by an analysis of the distribution of tubulin in the synapse, and of its amounts, in relation to the SV density (Wilhelm et al., 2014). Tubulin density enriches non-linearly with the SV density, suggesting that this protein indeed preferentially locates to SV clusters. However, other scenarios could also be imagined, under different synapse function conditions, with microtubule polymerization creating tubulin sinks away from the synapse, where limited levels of soluble tubulin remain.

To investigate whether SVs can bind soluble tubulin, we first considered the distribution of soluble tubulin in pre-synapses, by reanalyzing our data from (Reshetniak *et al*., 2020) and then experimentally investigating the interaction of SVs, isolated from rodent brains, with purified tubulin, by different imaging analyses. We found that the SVs were able to collect tubulin, confirming the previous FRAP-derived observations, and suggesting that SV-induced tubulin accumulation in synapses may be an important parameter for both synaptic and microtubule dynamics.

## Materials and Methods

### Analysis of soluble tubulin distribution in synapses

Experimental data was retrieved from (Reshetniak *et al*., 2020). In short, protein mobility data was collected using FRAP on synapses of cultured neurons, while electron microscopy provided detailed synaptic morphology reconstruction. A 3D synapse model was constructed, where particle movement was simulated by modeling particle trajectories, incorporating parameters as diffusion speed and SV binding, to reproduce the experimental fluorescence recovery recordings. The generated individual protein trajectories were then combined with quantitative data on protein abundancies from (Wilhelm *et al*., 2014), to generate detailed map of protein distribution in synapses. For more details on data acquisition and modeling, refer to the original work (Reshetniak *et al*., 2020).

Using the previously generated data on protein abundancies, positions, and synaptic geometry, we then calculated the local concentrations of various soluble proteins in axons, SV-free synaptic space, and within the SV cluster, which we define as the space occupied by the SVs and their immediate surrounding of twice their size. Using the obtained number we then calculated the enrichment of respective proteins in the SV cluster, as reported in Fig. 1.

**Figure 1.**
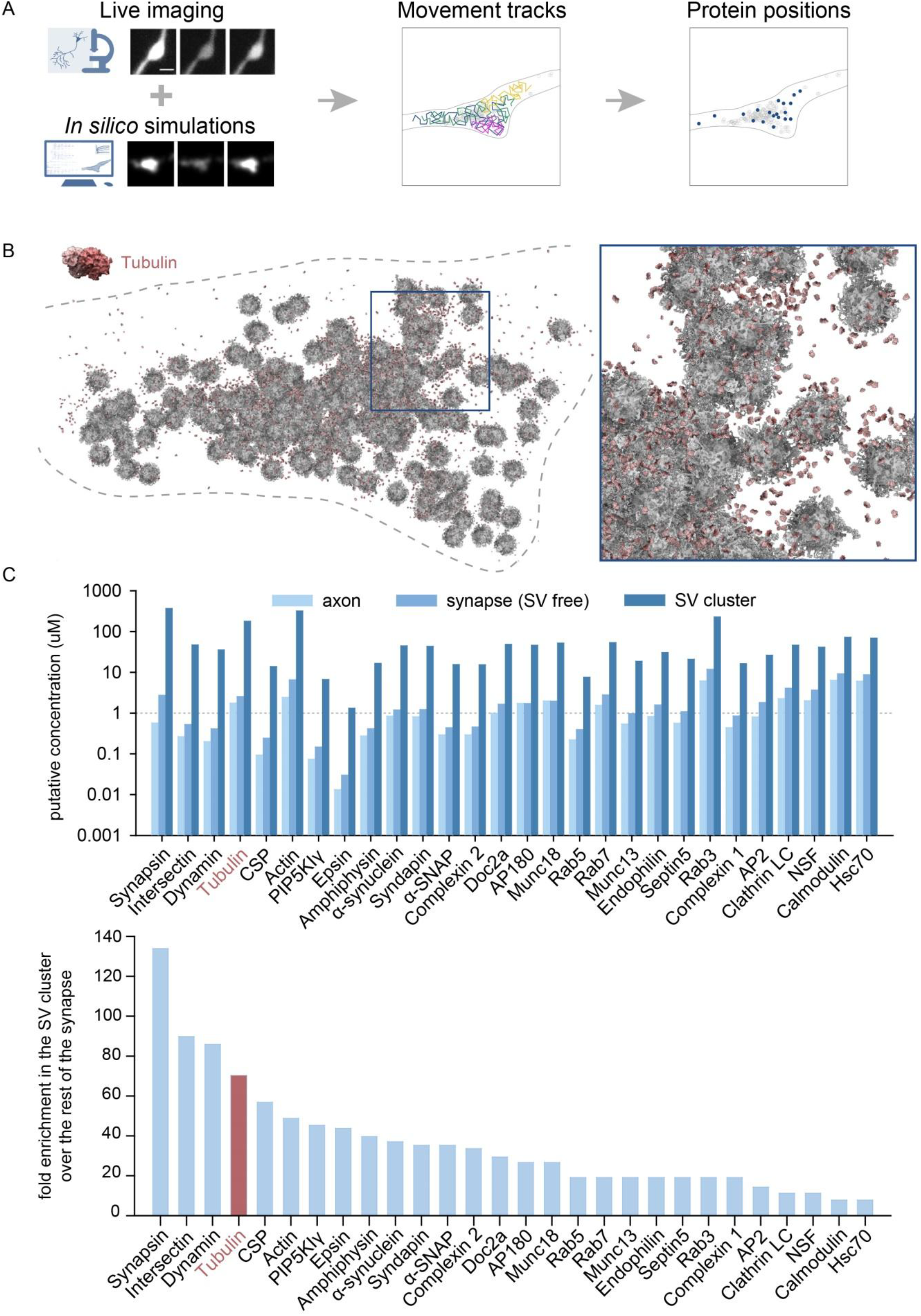
Tubulin distribution in the synaptic bouton. A. Schematic explanation of the experimental workflow of (Reshetniak *et al*., 2020). Live imaging was combined with *in silico* simulations to generate individual protein movement tracks, from which the positions of molecules could be extracted. **B**. Visualization of soluble tubulin dimers in a synaptic bouton, based on results from (Reshetniak *et al*., 2020). Dashed line Inset: a zoom-in into the synaptic vesicle cluster, demonstrating a higher local density of tubulin molecules. **C**. Quantification of the concentration of various soluble proteins in axons, SV-free synaptic space, and the SV cluster (top) and of the molar enrichment of respective proteins in the SV cluster, compared to the synaptic space outside of the cluster. Tubulin is among the most enriched proteins.

### Synaptic vesicle purification

Murine SVs were purified using the modified protocol from (Nagy *et al*., 1976; Huttner *et al*., 1983; Ahmed *et al*., 2013; Hoffmann *et al*., 2021). In brief, adult C57BL/6J mice (1:2 male:female ratio) were sacrificed and their whole brains including cerebellum extracted and homogenized in the homogenization buffer: 320mM Sucrose, 5mM HEPES-NaOH, pH 7.3, supplemented with 0.2 mM PMSF and 1 mg/mL Pepstatin A. Cellular debris was removed by centrifugation at 900 g, while supernatant was further centrifuged at 12,000 g. The white synaptosome-containing pellet was carefully washed with Homogenization buffer and centrifuged at 14,500 g. The obtained pellet was subjected to a hypo-osmotic lysis in the lysis buffer: 5 mM HEPES-KOH, 0.2 mM PMSF, 1 mg/mL Pepstatin A, pH 7.3. Lysate was centrifuged at 32,500 g to pellet the contaminants and SV-containing supernatant was further ultracentrifuged at 247,000 g. Pellet was resuspended in 40 mM sucrose and fractionated on a continuous sucrose density gradient (50–800 mM sucrose) by ultracentrifugation in SW 40 Ti rotor (Beckman) at 29,500 RPM for 3 h. All sucrose solutions were buffered with 5 mM HEPES-KOH, pH 7.3. SV-containing fraction was collected from the gradient, pelleted by centrifugation at 247,000 g, resuspended in the final reaction buffer: 150 mM NaCl, 25 mM Tris-HCl, 0.5 mM TCEP, pH 7.4 after which it was snap-frozen in liquid nitrogen and stored at -80°C until use.

The quality control of SVs was validated by Western Blot. Fractions were taken from SV prep brain homogenate, supernatant and pellet after lysate centrifugation and final SVs. All fractions were resolved in 12% Tris-Glycine SDS-PAGE gel and transferred onto Amersham Hybond LFP membrane (0.2 μm pore size, Cytiva). Wet transfer was done overnight at 10 V and 37°C. Membrane was washed in TBS and blocked in 5% milk or 3% BSA in TBS (depending on the primary antibody combination) for 1 h at room temperature. Incubation with primary antibodies was done overnight at 4°C with gentle shaking. Antibodies used: VAMP2 (SySy 104211 1:8000 in 3% BSA/TBS), Synaptophysin 1 (SySy 101011 1:1000 in 3% BSA/TBS), SDHA (EMD Millipore MABN630 1:1000 in 5% milk/TBS), PSD95 (SySy 124011 1:1500 in 5% milk/TBS). Staining with HRP-conjugated secondary antibodies (anti-mouse Sigma A4416 1:5000 in 5% milk/TBS or 3% BSA/TBS depending on primary antibody combination) was performed for 1 h at room temperature. Final blot was developed using Amersham ECL Prime Western Blotting Detection Reagents (Cytiva) and imaged on Fusion FX device (Vilber).

### Endosome purification

Endosomes of HEK293 cells were fluorescently labeled by uptake of Alexa Fluor 594-Dextran and isolated as described previously (Rizzoli *et al*., 2006).

### Liposome preparation

The following lipids were purchased from Avanti Polar Lipids: POPC (1-palmitoyl-2-oleoyl-sn-glycero-3-phosphocholine), POPE (1-palmitoyl-2-oleoyl-sn-glycero-3-phosphoethanolamine), POPS (1-palmitoyl-2-oleoyl-sn-glycero-3-phospho -L-serine), bovine liver polar lipid extract, and cholesterol. Texas Red DHPE (Texas Red 1,2-Dihexadecanoyl-sn-glycero-3-phosphoethanolamine) was purchased from Thermo Fisher. Large unilamellar liposomes were prepared by reverse-phase evaporation as described (Schmidt *et al*., 2020). For the chemically defined (CD) mix, lipids were mixed in chloroform at a molar ratio of 49.8:25:10:15:0.2 (POPC:POPE:POPS:cholesterol:Texas Red DHPE). Liver polar lipid extract (LPLE) was supplemented with an approximately matching amount of Texas Red DHPE. Chloroform was subsequently removed using a rotary evaporator by lowering the pressure step-wise to 20 mbar. The lipid film was then dissolved in diethyl ether to a final concentration of 20 mM. 300 μL of buffer (20 mM HEPES/KOH pH 7.4, 150 mM potassium chloride) was added, and the sample was sonicated for 1 min on ice (Branson Sonifier 450, 100% duty cycle, microtip limit 1). Afterwards the ether was removed at 500 mbar. After 10 min, additional 700 μL of buffer was added and the pressure was gradually decreased to 100 mbar until diethyl ether was completely removed. The resulting lipid suspension was extruded through a polycarbonate filter (11 x through a 0.4 μM filter, 21x through a 0.1 μM filter) using the mini extruder kit (Avanti Polar Lipids).

### SV-tubulin *in vitro* interaction assay

Circular 18 mm No. 1.5 glass coverslips were coated with BSA by incubation in 5% BSA in PBS overnight. 1 fmol of SVs per coverslip were added and centrifuged for 1 h at 4 000 g to force SV adhesion to the coverslips. Supernatant with unbound SVs was removed and remaining SVs were stained with 25 nM FluoTag®-X2 anti-Synaptotagmin 1 coupled to Abberior Star 635P (NanoTag Biotechnologies, Göttingen, Germany) for 30 minutes in a humidifying chamber at room temperature. The coverslips were then washed with PBS containing 2.5% BSA, followed by a wash with General Tubulin Buffer (80 mM PIPES pH 6.9, 2 mM MgCl_2_, 0.5 mM EGTA) (Cytoskeleton, Inc, Denver, USA) supplemented with 10% glycerol and 1 mM GTP. The coverslips were then incubated with 5 mg/ml tubulin solution, consisting of 95% unlabeled porcine brain tubulin (Cytoskeleton, Inc, Denver, USA) and 5% fluorescently labeled (HiLyte Fluor™ 488) tubulin (Cytoskeleton, Inc, Denver, USA) in General Tubulin Buffer supplemented with 10% glycerol and 1 mM GTP for 30 minutes at 37°C. Following the incubation, unbound tubulin was washed off using stabilization buffer (General Tubulin Buffer supplemented with 30% glycerol, 1 mM GTP, and 5 μM paclitaxel) and any microtubules formed on the SV were stabilized by incubation in stabilization buffer for 5 minutes at 37°C. The samples were then fixed with 4% PFA in PBS for 20 minutes, quenched with 100 mM NH_4_Cl in PBS for 20 minutes and mounted in Mowiol for further microscopic evaluation.

In parallel, control experiments were performed following the same protocol, but in the absence of the SVs, with fluorescently labeled endosomes instead of the SVs, or using non-specifically binding antibodies.

### Microscopy and image processing

Widefield images were obtained using an inverted Nikon Ti-E epifluorescence microscope (Nikon Corporation, Chiyoda, Tokyo, Japan) equipped with a 60 × 1.4 NA oil-immersion Plan Apochromat objective (Nikon Corporation, Chiyoda, Tokyo, Japan), an HBO-100W Lamp and an IXON X3897 Andor (Belfast, Northern Ireland, UK) camera.

Confocal images were obtained using a TCS SP5 microscope (Leica, Wetzlar, Germany) equipped with a HCX Plan Apochromat 100× 1.4 NA oil-immersion objective. The 488 nm line of an Argon laser was used for excitation of HiLyte Fluor™ 488, while Alexa Fluor 594 and Abberior Star 635P were excited using HeNe 594 nm and HeNe 633 nm lasers, respectively. Images were collected at 512×512 pixel resolution and physical size of 62×62 μm. Acquisition settings were kept identical across experiments.

Exemplary images were processed in ImageJ 1.54f (NIH, USA). For quantification, custom-written MATLAB (The MathWorks Inc, Natick, MA, USA) routines were employed. The images were subjected to an empirically-defined, automated thresholding procedure, to isolate tubulin-positive regions. The center-of-mass of the respective regions was determined automatically, in the tubulin channel, and line scans were performed automatically, through the pixel corresponding to the center of mass. The peak intensity was measured for tubulin and for the other channels, from the respective line scans. The intensities observed were used to classify the TRRs into SV-associated or non-associated classes, according to the synaptotagmin signals, and to measure the non-specific signals associated to the tubulin spots.

## Results

Our own observations of soluble, GFP-tagged tubulin (Reshetniak *et al*., 2020), expressed in cultured hippocampal neurons, suggest that abundant levels of soluble tubulin are present in the synapses. Using the data from the mentioned study, we performed further analysis of the distribution of soluble tubulin in the synaptic bouton, which demonstrated higher local density of tubulin molecules in the vicinity of the SVs (Fig. 1A, B). Indeed, calculations of the molar enrichment of various proteins in the synapse suggests that tubulin enriches strongly in the SV cluster, being the 4^th^-most enriched molecule in this environment (**Fig. 1C**). This is an unexpected observation, since the magnitude of the enrichment of tubulin in the SV cluster is substantially higher than for *bona fide* SV-binding molecules, as Rab3, or for vesicle cluster components, as alpha-synuclein (**Fig. 1C**).

This computational analysis suggests that SVs might accumulate soluble tubulin on their surface. To directly observe this possible interaction of soluble tubulin with SVs, we relied on a simple setup, as illustrated in **Fig. 2**. We immobilized purified brain-derived SVs (from the brains of adult mice) on glass coverslips and incubated them with a solution of purified tubulin (5% fluorescently labeled). We then analyzed the distribution of tubulin, relative to the SVs, by fluorescence microscopy.

**Figure 2.**
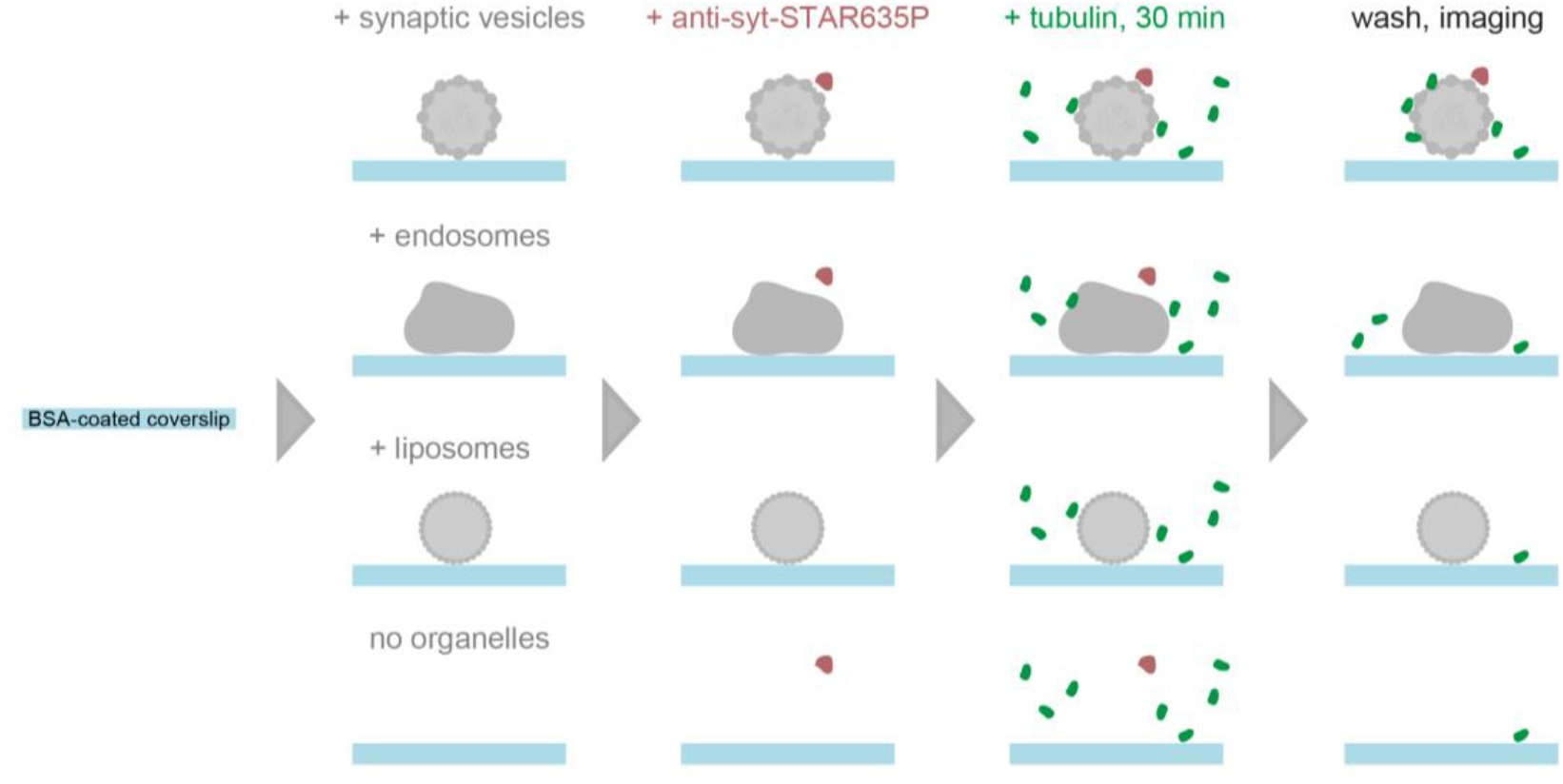
Scheme of the experimental workflow. Synaptic vesicles are allowed to adhere onto BSA-coated glass coverslips, and are immunostained with fluorescently-labeled anti-synaptotagmin nanobodies. A buffer containing native and fluorescently labeled tubulin is then added, and is incubated for 30 minutes at 37°C. All unbound proteins are then washed off, and the samples are analyzed by fluorescence imaging. The same procedure is performed in the presence of fluorescently labeled endosomes, liposomes, or in the absence of any organelles, as controls.

We found that soluble tubulin tends to accumulate in distinct loci, which we term tubulin-reach regions (TRRs). These TTRs often, but not always, colocalize with the clusters of SVs, and are also formed in the absence of the SVs (**Fig. 3A, Fig. S1A**,), implying that two populations of TRRs form, SV-associated and SV-non-associated ones. Average line scans through automatically selected synaptotagmin-positive loci indicate a clear overlap between the SVs and SV-associated TRRs (**Fig. 3B, C, Fig. S1B**).

**Figure 3.**
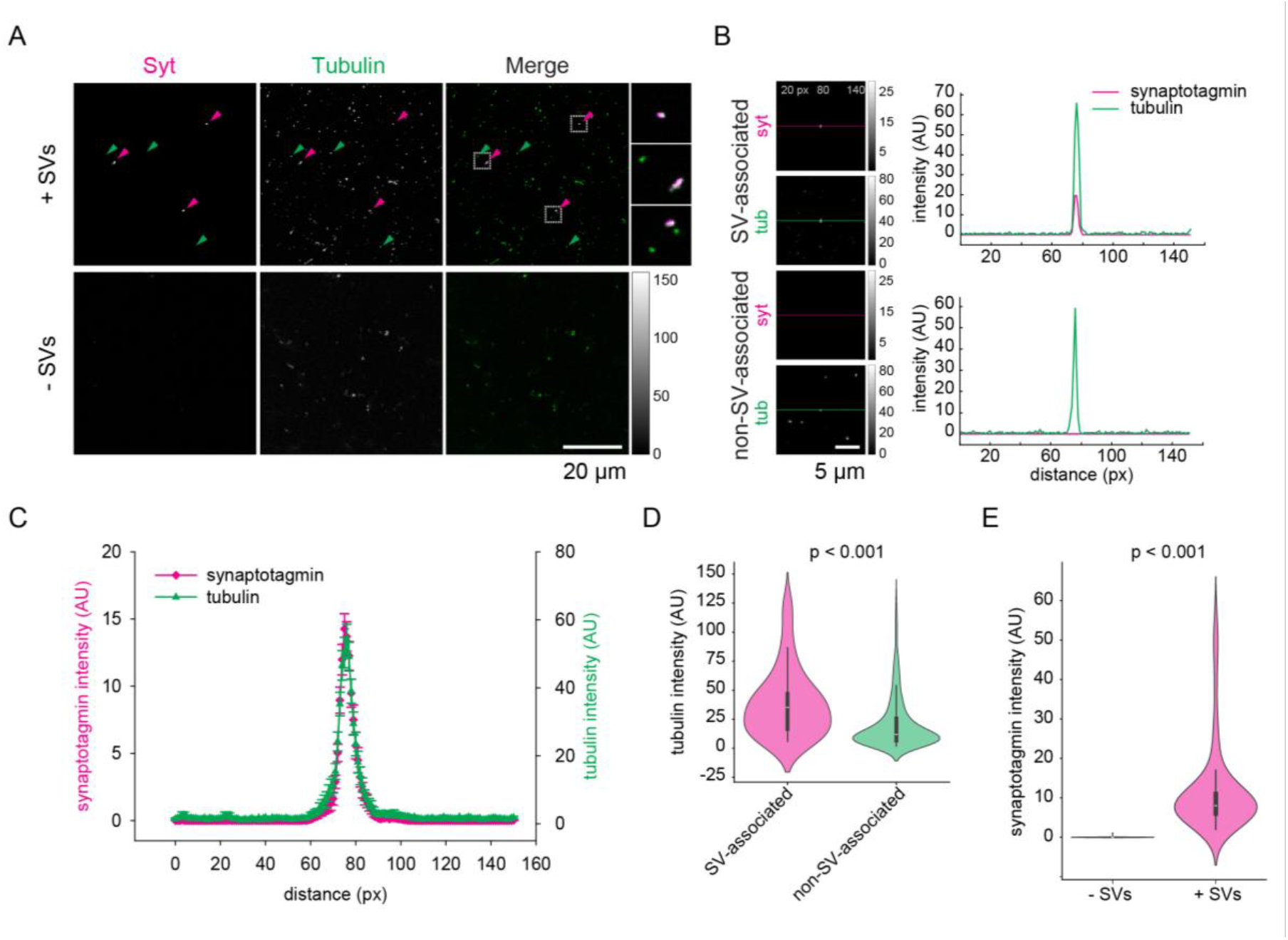
Tubulin accumulates in SV-enriched areas. A. Confocal view of tubulin distribution in the presence and in the absence of SVs. Arrowheads indicate SV-associated (magenta) and non-SV-associated TRRs (green). Zoom-ins into the squared regions are shown on the right. **B**. Left: exemplary images with automatically-detected TRRs in the center and corresponding synaptotagmin signal. Colored lines indicate the position where line scans are taken 75 pixes in both directions from the center of the identified spot. TRR are classified into SV-associated (top) or non-SV-associated (bottom) based on the intensity of the synaptotagmin signal in the region masked by tubulin channel. Right: line scans from the regions showed on the left. **C**. Average line scans indicating tubulin signals overlapping with SVs, as detected in confocal imaging. **D**. Comparison of tubulin intensity in TRRs associated or not associated with SVs. Data from 5 independent experiments, 55 SV-associated TRRs and 540 non-SV-associated TRRs were quantified. Indicated significance bracket: Kruskal-Wallis test, p = 0.0008. **E**. Synaptotagmin intensity in samples with or without SVs. Data from 3 independent experiments, 89 TRRs from samples containing SVs and from samples not containing SVs were quantified. Wilcoxon rank sum test, p = 9.8111×10^−34^.

SV-associated TRRs had a higher intensity than TRRs forming in the same samples, but in SV-free areas (**Fig. 3D, Fig. S1C**). The intensity of the latter was also lower than that of TRRs forming in the absence of SVs (in samples containing only tubulin, but no other molecules), as indicated in **Fig. S1C**. This observation suggests that SVs sequester tubulin, reducing the availability of these molecules for the formation of TRRs in other regions. To validate the SV interaction to the TRRs, we quantified the intensity of anti-synaptotagmin staining in the presence and in the absence of SVs. The results suggest that the colocalization observed between TRRs and SVs is not affected by autofluorescence or other artefacts (**Fig. 3E, Fig. S1D**).

Notably, in both widefield and confocal images, we only sporadically observed short microtubules, suggesting that the interaction of SVs and tubulin may not automatically translate to microtubule polymerization, in spite of the presence of paclitaxel, which should stabilize newly formed microtubules.

In principle, these observations could be explained by the non-specific interaction of tubulin with any membranes provided, or by the binding of both tubulin and SVs to adherent patches on the coverslips. To test the first of these hypotheses, we repeated the experiments in the presence of fluorescently labeled endosomes, isolated from HEK293 cells, or artificial liposomes of various compositions instead of SVs. As shown in **Fig. 4A**, tubulin demonstrated seemingly random distribution, with no clear preferential localization close to endosomes or liposomes. The intensity correlation analysis indicated virtually no colocalisation with liposomes of either chemically-defined (CD) composition or derived from a liver polar lipid extract (LPLE), and moderate, but considerably less pronounced compared to the SVs, colocalization with cell-derived endosomes (**Fig. 4B, S2**). The significantly increased amount of tubulin retained in the endosome-containing sample can be explained by its binding to other cellular components present in the endosomal preparation, which are, however, not fluorescently labeled.

**Figure 4.**
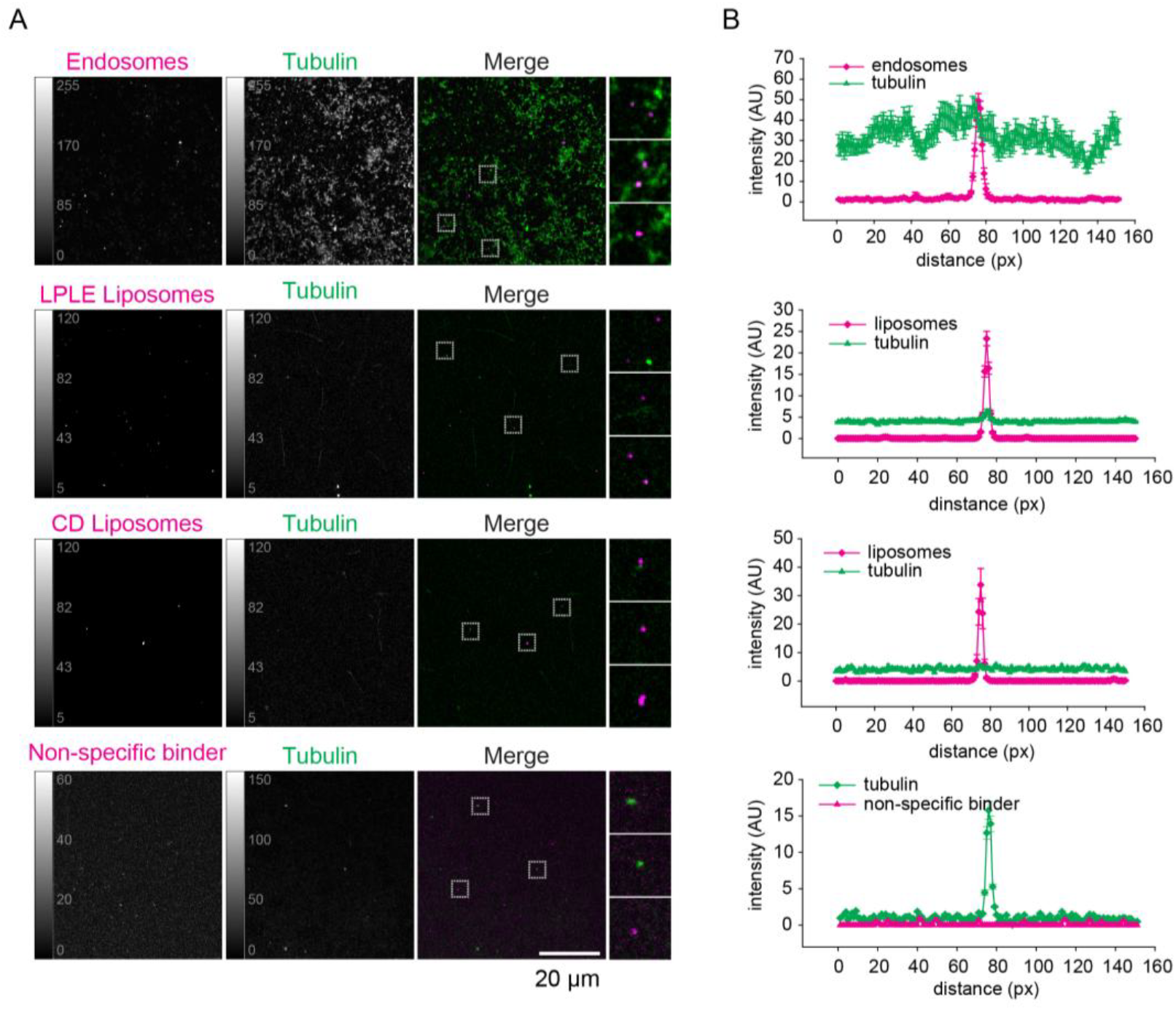
Tubulin does not preferentially accumulate on endosomes, artificial vesicles, or at sites of non-specific protein adhesion. A. Confocal views of tubulin distribution in samples where fluorescently labeled endosomes, two types of liposomes (LPLE – liver polar lipid extract, CD – chemically defined composition, 49.8:25:10:15:0.2 POPC:POPE:POPS:cholesterol:Texas Red DHPE), or non-specific binder (anti-rabbit antibody, used with no rabbit antibodies in the sample) were attached to the coverslips instead of synaptic vesicles. **B**. Averaged line scans from the corresponding samples. Data from 3 independent experiments, 30 endosome-positive regions, 178 LPLE-liposome-positive regions, 25 CD-liposome-positive regions, 86 tubulin-positive regions for the non-specific stainings.

To test the second hypothesis, of the possible coincidental binding of both tubulin and SVs to adherent patches on the coverslips, we incubated the coverslips with tubulin and a fluorescently labeled antibody, in the absence of an antigen, to reveal possible locations of non-specific binding on the coverslip surface. An analysis of the relative distribution of TRRs and the non-specific binder revealed effectively zero overlap between these signals (**Fig. 4B, S2**).

Apart from the non-specific adherence of tubulin to the coverslips, the observed signal accumulations might be mediated by fluorophore interactions. To test this, we repeated the experiments using a lower tubulin labeling ratio, while preserving the same total tubulin concentration. This resulted in preserved colocalisation, but decreased tubulin signal intensity (**Fig. S3**), indicating that non-labeled tubulin outcompetes the labeled molecules, therefore confirming the role of tubulin, not the fluorophore, in establishing this interaction. Lastly, we tested whether visualizing SVs by targeting another SV protein will affect the relative distribution of SV and tubulin signal. We repeated the experiments, immunostaining for synaptophysin, which resulted in tubulin accumulation around synaptophysin-positive regions (**Fig. S3**).Taken together, these experiments indicate that the colocalization of SVs and tubulin is caused by a specific interaction between them.

## Discussion

Overall, these experiments indicate that isolated SVs are able to collect soluble tubulin. This observation fits well with the high enrichment of tubulin within the SV clusters, as estimated from previous imaging experiments in hippocampal cultured neurons (Reshetniak *et al*., 2020).

The current view of the SV cluster organization is that the SVs, along with at least some of the synapse-resident soluble proteins, participate in liquid-liquid phase separation (LLPS; (Milovanovic & Camilli, 2017; Reshetniak *et al*., 2024)). We have previously analyzed the motion of 28 soluble proteins in the synapse, by FRAP and particle tracking with electron microscopy, nanoscale imaging and modeling. We could demonstrate that soluble protein mobility is strongly affected by interactions with the SV cluster, and that the organization of many of these proteins can be, indeed, approximated by a liquid-liquid phase separation between the vesicle cluster and the rest of the presynaptic bouton (Reshetniak *et al*., 2020).

The most enriched proteins in the vesicle cluster were synapsin and intersectin, which have been shown to phase-separate with lipid vesicles *in vitro* (Milovanovic *et al*., 2018). The third most-enriched protein was dynamin, which, like synapsin, is an SH3-domain-binding protein and also contains a large intrinsically disordered region, making it probable that it can participate in the formation of such a liquid phase. Assuming that these molecules are necessary and sufficient for the formation of the synapse-resident LLPS phase (Milovanovic *et al*., 2018; Sansevrino *et al*., 2023; Hoffmann *et al*., 2023; Alfken *et al*., 2024), it is evident why such a phase forms only in boutons, but not along the axon: the synapsin, intersectin and dynamin concentrations are far too low in the axon, when compared to the levels needed *in vitro* for liquid phase formation.

It is still unclear, however, whether other soluble proteins (other than synapsin, intersectin and dynamin) also participate in a liquid phase-like arrangement in the synapse. The 5^th^-most enriched protein in the SV cluster is CSPalpha, a chaperone that is strongly associated to the SV membrane (Gorenberg & Chandra, 2017), and does not seem to unbind from the vesicles during synaptic function, even over long time periods (Truckenbrodt *et al*., 2018). The 6^th^-most enriched protein is actin, whose association with synapsin and SVs has long been demonstrated (Cesca *et al*., 2010). Therefore, the most enriched proteins in the SV cluster are either SV components (CSPalpha), proteins with large intrinsically disordered domains, which are expected to participate in LLPS (synapsin, intersectin, dynamin), or their interactors (actin).

The obvious exception is tubulin, which is not classical LLPS-forming molecule. Its recruitment to SVs may be due to direct interactions with SV membrane components (as synaptotagmin), or to interactions with the abundant, and LLPS-inducing, synapsin molecules, as discussed in the Introduction. As these molecules are found to levels of ∼8 copies per purified SV (Takamori *et al*., 2006), they would certainly be able to recruit tubulin.

Would recruited tubulin then participate in LLPS dynamics in the synapse? Recent observations suggest that tubulin is able to phase-separate *in vitro*, if suitable interactors are present. For example, Abl2, a member of the Abl kinase family, is able to undergo LLPS separation (induced by its C-terminal half), and forms co-condensates with tubulin (Duan *et al*., 2023). Moreover, this process appears to concentrate tubulin locally, to promote microtubule nucleation, which may be especially important for microtubule repair (Duan *et al*., 2023). Other examples include the end-binding protein EB1, which forms condensates that recruit tubulin dimers, along with other plus-end tracking proteins (Song *et al*., 2023), or calmodulin-regulated spectrin-associated protein 2 (CAMSAP2), which also co-condensates with tubulin (Imasaki *et al*., 2022).

Nonetheless, it remains unclear whether tubulin participates in synaptic LLPS (Fig. 5). Neither FRAP experiments, nor common LLPS-manipulating drugs, like 1,6-hexanediol, can provide a clear answer. This is because the behavior of the molecules could be explained both by bona fide LLPS arrangements and by binding and unbinding from low-mobility objects, such as SVs. In living cells, this ambiguity is difficult to resolve (Muzzopappa *et al*., 2022). Moreover, tubulin may exhibit complex dynamics in synapses, separating into freely soluble, SV-bound and polymerized pools of molecules. This behavior will not be easily tracked in either FRAP or 1,6-hexanediol treatments.

**Fig. 5.**
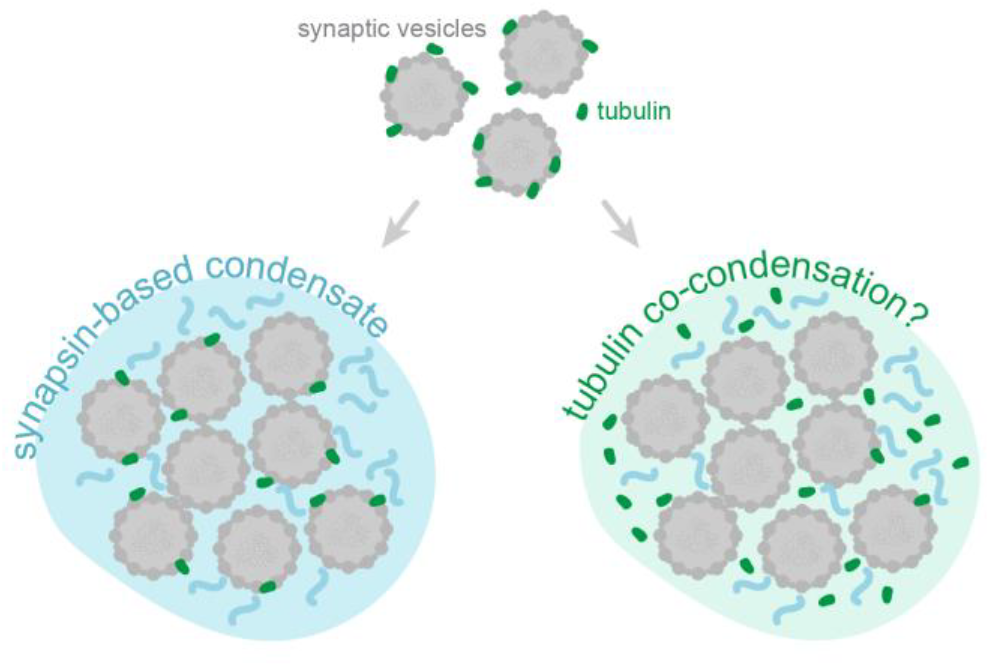
Possible organizarion of tubulin in the context of presynaptic LLPS. Our data demonstrates, that soluble tubulin can bind free synaptic vesicles. Tubulin can be then sequestered to the SVC through these direct interactions with the SVs. Alternatively, it might undergo phase transitions together with other phase-separating proteins of the SVC.

In our present experiments, the SVs are immobilized, preventing them from movement, which is a pre-condition of condensate formation. This implies that our experiments do not provide conditions under which LLPS processes could be tested. We suggest that dedicated experiments should be performed, *in vitro*, in conditions enabling both SV and tubulin movement. This would allow to test whether they indeed co-condensate. In fact, our observations are also compatible with the hypothesis that SV clusters act as binding sites for free tubulin, thereby sequestering these molecules, and preventing microtubule polymerization within the clusters. This idea is in line with our observations that the SV-bound tubulin virtually never generates microtubules, even in the presence of taxol, which should promote microtubule formation.

Overall, it is quite likely that LLPS formation in the SV cluster takes place in parallel with tubulin sequestration, without tubulin actively participating in LLPS processes.

### Limitations of this study

The experiments reported here were performed in a simplified *in vitro* system, with typical limitations of such approaches. First, they do not consider contributions of other cellular components that might modulate SV-tubulin interactions. Tubulin concentration and SV density do not recapitulate such of intrasynaptic ones, which might also have an effect on the dynamics of their interaction. Additionally, the mixed sourcing of the SVs, endosomes, and tubulin might affect the interactions due to differences in e.g. post-translational modifications between model systems. Lastly, our SV purification protocol does not discriminate between neurotransmitter carrying vesicle type, meaning that it is impossible to state whether tubulin-binding ability of SVs is limited to synapses of a specific type, based on the results presented here. The current work represents a proof-of-concept study and further investigations of SV-tubulin interactions focused on their analysis in native conditions of a synapse are warranted.

## Author contributions

SR and SOR conceived the study, TM and AK performed the experiments, AS provided the liposomes, DM, SOR, SK, and SR supervised the study, SOR and SR analyzed the data, DM, SOR and SK acquired funding, SR and SOR wrote the initial draft of the manuscript, all authors refined the final manuscript.

