## Supplementary figures for "Brain-derived synaptic vesicles have an intrinsic ability to sequester tubulin"

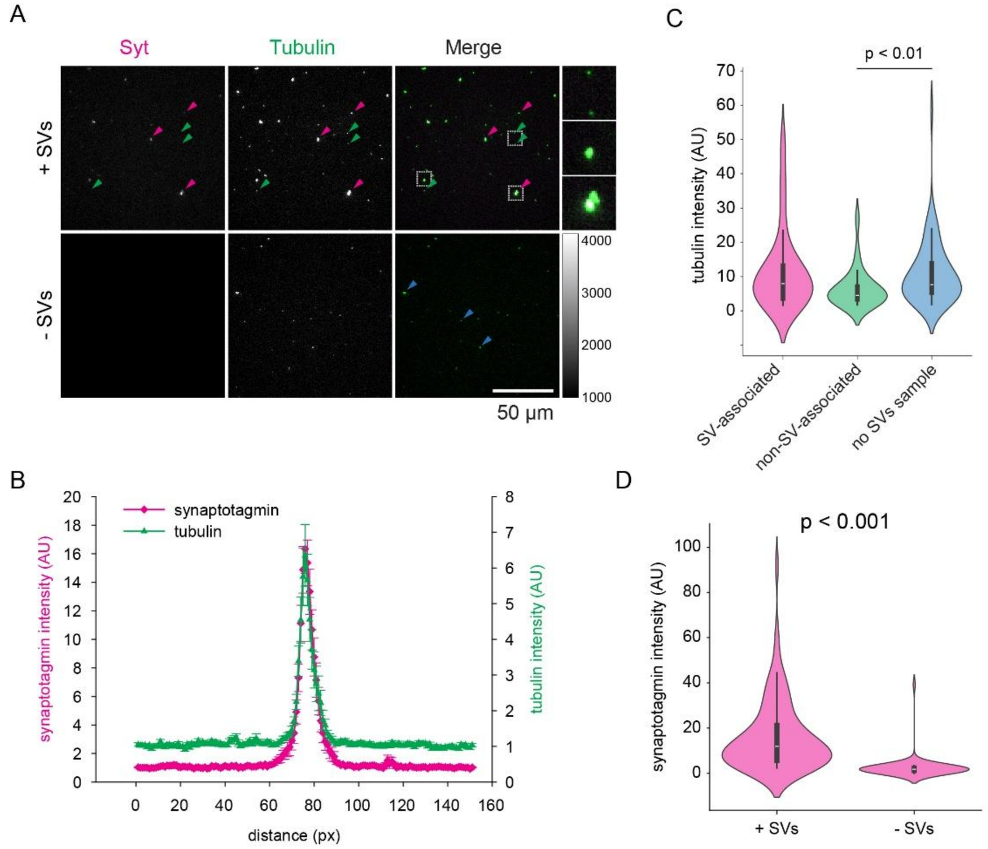


**Fig. S1. Tubulin accumulates in SV-enriched areas. A.** Widefield view of tubulin distribution in the presence and in the absence of synaptic vesicles. Colored arrowheads indicate examples of different tubulin-rich regions (TRRs), which are analysed in panel C. **B.** Average line scans of tubulin signals overlapping with SVs. **C.** Comparison of tubulin intensity in different TRRs, based on their association with the SVs, as determined by the presence of the anti-synaptotagmin signal. Refer to panel A for examples: magenta arrowheads indicate TRRs associated with SVs, green arrowheads – TRRs not associated with SVs in a sample containing SVs, and blue – TRRs observed in the samples without SVs. Data from 3 independent experiments, 48 SV-associated TRRs, 27 SV-non-associated, and 60 TRRs in a sample lacking SVs were quantified. Indicated significance bracket: Kruskal-Wallis test, $p = 0.0087$. **D.** Synaptotagmin intensity in TRRs in samples with or without SVs. Data from 3 independent experiments, 86 regions from samples containing SVs and 60 regions from samples not containing SVs were quantified. Wilcoxon rank sum test, $p = 4.888\times10^{-21}$.

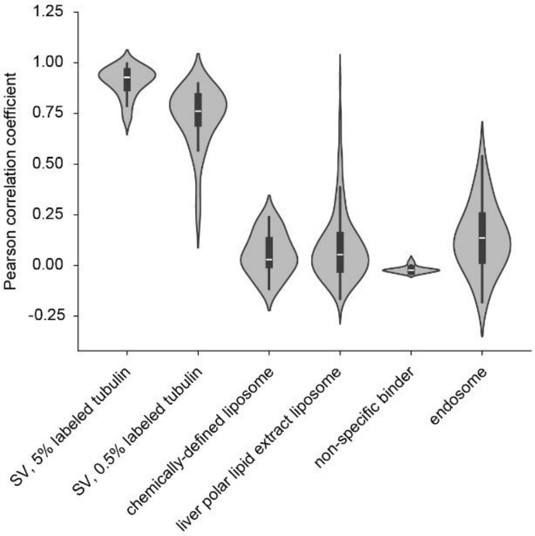


**Figure S2. Pearson correlation coefficients of tubulin and various organelles' intensity.** Each data point represents the Pearson R value, calculated for the two linescans in the tubulin and organelle images.

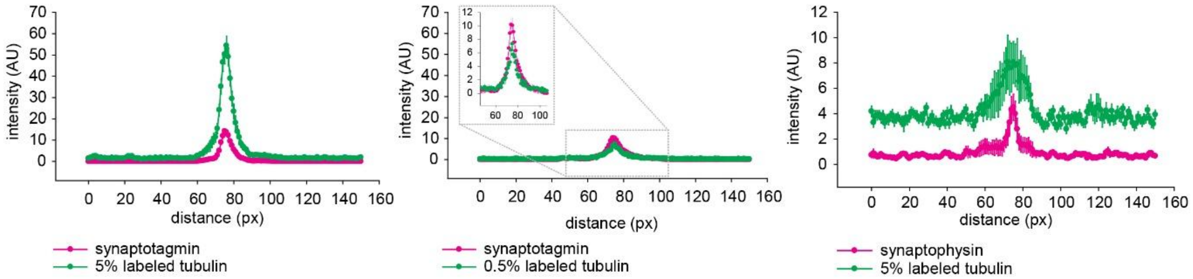


**Figure S3. Comparison of average line profiles of tubulin and SV marker intensities of TRRs detected in samples using different labeling approaches.** Decreasing the fraction of fluorescently labeled tubulin or targeting another SV protein does not affect the relative distribution of SVs and tubulin.
